# Anterior insular cortex firing links initial and sustained encoding during aversion-resistant alcohol consumption

**DOI:** 10.1101/2022.05.24.493243

**Authors:** Phillip Starski, Mitch Morningstar, Simon Katner, Raizel Frasier, Thatiane De Oliveira Sergio, Sarah Wean, Christopher Lapish, F Woodward Hopf

## Abstract

Compulsive-like alcohol drinking (CLAD), where intake persists despite adverse consequences, is often a core facet of alcohol use disorder. Recent work sheds light on underlying mechanisms, but much remains unknown about CLAD etiology. Previously, we showed that projections from anterior insula (aINS), a central mediator of emotion, motivation, and interoception, promote CLAD in rodents, and heavy human drinkers exhibit similar insula-circuit recruitment under compulsion-like conditions. However, global aINS inhibition also reduces alcohol-only drinking (AOD), and one major obstacle is the lack of information on aINS firing patterns that could promote different aspects of intake. Here, we recorded single-unit activity in right aINS from 15 rats during AOD or CLAD (10mg/L or 60mg/L quinine in alcohol). Neurons with a sustained-increase or sustained-decrease phenotype (SIP, SDP) showed no firing differences across drinking conditions. In contrast, aINS neurons with a phenotype of strong firing increase at initiation of responding (IRP) showed significantly greater activity across the rest of licking during CLAD versus AOD, concurring with our previous behavioral findings suggesting quick evaluation and response strategy adjustment under CLAD. There were also no condition-related differences in firing-phenotype abundance. Further, total responding only correlated with abundance of SDP cells, but SDP firing returned to baseline during pauses in licking, while IRP and SIP sustained responding through pauses in licking. Thus, only aINS cells with a particular strong firing at licking onset showd greater sustained responding under compulsion-like conditions, while other cells likely promoted drinking more generally, providing critical new information about how aINS activity could promote alcohol consumption under different drinking conditions.

## INTRODUCTION

Alcohol Use Disorder (AUD) is among the most prevalent mental disorders [1,2], which leads to ~$250 billion in avoidable costs per year in the US alone, and is associated with many economic, social, and medical harms [3–10]. Also, while decreasing alcohol intake reduces risk of harm and relapse [8–14], there are limited treatment options for AUD [15], representing an area of critical unmet need.

Among factors that can contribute to problem drinking, compulsion-like alcohol drinking (CLAD), defined as continued intake despite negative consequences, can strongly promote intake and thus be a major obstacle for AUD treatment [16–25]. CLAD-like behaviors feature prominently in the DSM-V description of AUD (e.g. [14,26]), and greater drinking is related to more alcohol problems [11–14] (addressed further in Discussion). Thus, to improve treatment, it is critical to uncover mechanisms that underlie aspects of problem drinking, including aversion-resistant consumption patterns. In order to better understand pathological intake mechanisms like CLAD, rodent models can be of value, since willingness to respond for reward paired with adverse consequences is considered to model some aspects of compulsion-like responding in humans [20,21,24,25,27].

Alcohol drinking is likely driven by its salient motivational properties, making anterior insula (aINS) a probable contributor. The aINS, a key regulator of the Salience Network, mediates rapid identification and responding to important situations [28–32], and human aINS is linked to many aspects of alcohol drinking [28] (see Discussion). Importantly, we have previously shown that specific aINS projections, aINS-Nucleus Accumbens Core (NAcb) [33] and aINS-Locus Coeruleus area [34], are essential for promoting CLAD in male rodents, with little impact on alcohol-only drinking (AOD), suggesting particular aINS importance for CLAD (see also [35,36]). In interesting concurrence, compulsion-like responding for alcohol in human AUD involves activation of a similar aINS-striatal circuits [37,38], and risk- and aversion-related responding more generally relates to aINS activity (reviewed in [28]). However, global inhibition of aINS can strongly impact AOD as well as CLAD ([34,39], consistent with insula/salience network importance for motivated behavior more generally [28-30,32,40], and suggesting that there could be parallel aINS-related mechanisms that differentially regulate AOD and CLAD (discussed in [34]). However, one major obstacle to understanding aINS importance for promoting alcohol consumption is the lack of information on in vivo aINS firing patterns, which could reveal critical and foundational insights into the nature of aINS activity that drives different aspects alcohol intake.

Thus, here we used 32-electrode multi-wire electrophysiology to record single-unit activity in right aINS from 15 rats during different alcohol drinking sessions, including AOnly, what we have termed moderate-challenge CLAD (AQ10, alcohol with 10 mg/L quinine), and higher-challenge CLAD (AQ60, alcohol with 60 mg/L quinine) [27,33,34,41]. We used quinine-resistant drinking, rather than footshock, since similar cortical-NAcb projections underlie both quinine- and footshock-resistant alcohol consumption [33], and higher shock-resistance correlates with higher quinine-resistance for alcohol [42,43]. Together, these validate the use of the technically simple quinine-resistance for studying CLAD, and quinine-resistance is also easier to test in a graded manner to assess responding under different levels of challenge. In addition, when seeking to predict what aINS activity patterns might contribute to driving alcohol intake, we note that CLAD shows greater aINS cFos in mice than AOD [35], suggesting importance of higher aINS firing for CLAD (see also [36]). Further, aINS has been widely related to initiation of responding, consistent with Salience Network (above), and also linked to sustaining salience-driven behavior, with maintained aINS activity reflecting the main task goal and/or overall reward value of a situation [44–47]. Also, while there are quite limited studies in rodents, Guillem et al. (2010) [48] found that ~50% of aINS cells showed hours-long sustained plateau activity changes across a cocaine intake session (perhaps reflecting sustained drive for and overall value of cocaine responding) (see also [49]).

Thus, our findings link these predictions of greater activity and initiation- and maintenance-related firing for CLAD. Specifically, the set of neurons with strong firing at onset of licking (which we call Initial Response Phenotype, IRP) have significantly greater sustained activity across the remainder of licking for CLAD versus AOD. Linking initial with plateau firing during CLAD also concurs with our previous behavioral evidence that rats quickly determine what they are drinking (AOD or alcohol paired with aversion) and adjust their response strategy [27,41]. In particular, rats under CLAD have earlier and faster responding (what we have called the “Head Down and Push” model of CLAD, see Discussion), which predicts that CLAD involves a single, session-level action strategy which sustains conflict-resistant responding while minimizing the impact of adverse consequences. This is akin to “intake defense” [50], which assures that “sufficient” intake occurs under conditions of challenge (such as water deprivation or hunger), without titrating moment-to-moment responding to motivation level (see Discussion). This may explain why the number and firing level of cells with increased plateau firing (IRP and sustained increase phenotype, SIP) did not relate to actual alcohol consumption. In contrast, greater drinking levels overall were related to more cells with a phenotype of sustained decreases in firing (SDP) during intake, similar to some other work linking consumption to reduced neuronal firing (e.g. [51,52] but see [53,54]). However, firing in SDP cells returned to baseline during pauses in licking, while IRP and SIP licking-related firing changes were maintained during pauses in licking, suggesting that IRP and SIP contributed to the overall action strategy for alcohol consumption, while SDP cells were more related to actual licking. Taken together, we suggest that neurons with strong activity at onset of licking help evaluate the drinking condition, and greater sustained activity in these neurons then maintains drive for alcohol under adverse conditions. In parallel, neurons with decreased activity facilitate licking. Our work provides critical new insights into how aINS activity patterns could promote different aspects of alcohol drinking.

## METHODS

### Animals

Adult, male Wistar rats (Envigo, IN) arrived at ~PN55 and were single housed in clear plastic cages, with lights off 11am-11pm. After 2wk habituation to vivarium, alcohol drinking began, with experiments performed during their dark cycle under red light. All experiments were carried out in accordance with procedures approved by Indiana University Institutional Animal Care and Use Committee. Rats were ~450-750g at the time of studies.

### Alcohol Drinking Model

Rats initially drank under an intermittent access two-bottle-choice (20% alcohol in water, versus water-only in a second bottle) (IA2BC), with 16-24-hr intake each day starting Monday, Wednesday and Friday at ~30min into the dark cycle. After at least 3 months of IA2BC, rats began to drink under limited daily access (LDA), where they could drink for 20min/d, 5d/wk. Alcohol and water bottles were presented through holes in the front of the home cage ~7cm above the cage floor. After 3-4 weeks LDA, rats had 2-3 days/wk of drinking alcohol with quinine, usually 2 sessions each of AQ10 and AQ60, in part to habituate to quinine assure that alcohol-quinine is not novel during experimental sessions. We widely use these methods [27,33,34,41,55,56]. We note that IA2BC rats can drink ~0.8 g/kg in a 20-min session, with BECs of ~40-50 mg/dL [33,57]; we consider CLAD less about intake levels per se, and more defined by the motivational processes (i.e., the decision to drink alcohol despite aversive consequences).

### In vivo recording probes to assess global aINS firing patterns

In vivo recording methods follow those previously published from the lab of Dr. Christopher Lapish [58–61]. 32 channel Multi Wire Arrays were home-made (detailed in [61]), with 23 μm tungsten wires (California Fine Wire, CA) 100 μm apart threaded through a custom-made 16×2 silicon tube array (100 μm diameter silicon tubing, Polymicro Technologies, AZ). Wires were then attached to a custom-fabricated (developed by Likhtik Lab) electronic interface board (EIB) with gold pins. Intan headstages were connected to a male Omnetics connector that was soldered onto the EIB.

### Surgery

Animals underwent surgery with standard anesthesia (isoflurane) and analgesia (buprenorphine and carprofen for 2 days). Four stainless steel anchoring screws were strategically placed rostral and caudal to bregma, with two screws in the occipital bone (to attach grounding wires). A rectangular craniotomy was performed over aINS, the dura excised, and the site cleared of debris. The probe was descended into the target location (AP: +2.9, ML: +4.8, DV: −6.0 from bregma), cemented into place, and connected by wire to two ground screws in occipital bone. Animals recovered in a heated homecage until alert and mobile.

### Experimental Drinking Sessions

For experimental sessions, drinking time was reduced to 4 min per session, as we have done before [33,34], since rats strongly “front-load” alcohol in these brief sessions and have the vast majority of their intake in the first minutes of alcohol access (e.g. see examples in **Fig.2**). All experimental sessions occurred in the home cage. Rats were first habituated to drinking in the behavioral room, where wire cage lids were replaced by a cage extender, an in-house creation where the bottom of a home cage is removed, and placed upside down on top of the rat’s home cage and fixed in place. This “tower” allows access for head-attached cables but preventing rats from jumping out. Additionally, custom-created two-bottle holders were attached to the front of their homecage to accommodate angled sippers required for recording experiments.

Rats trained for ~2 weeks drinking in this tower and then underwent surgery to implant recording devices (above), followed by one week recovery. Then rats drank ~1 wk more, with daily handling, then ~1 week of drinking with dummy cable attached (as in [33,34]), before recording studies began. We then recorded in vivo aINS firing activities in AOD, AQ10, and AQ60 sessions tested within rat, with drinking conditions tested in randomized order. In addition, each drinking condition was tested at least twice within each rat, as we did in [27,33,34,41,55,56], with the goal of having two sessions from each rat for each drinking condition which had sufficient number of recorded cells and licking level in each session (detailed below).

Drinking patterns during experiments were assessed with custom-made capacitive lickometers, with custom-written C++ code running on an Arduino Uno with an Adafruit MPR121 capacitive touch sensing breakout board to record lickometry patterns. A very small capacitive current is run through the 7.6 cm metal licking tube, and the discharge is detected when the rat’s tongue contacts the metal tube. We note that this very small change in electrical charge, does not disrupt ability to record neurons: several groups have recorded in vivo firing during licking using very similar lickometers [62,63], including aINS firing during licking for tastants [64,65]. Also, we have performed extensive analyses using cross-correlograms looking for firing changes precisely timed with licks (which might suggest artifactual firing patterns), and saw little evidence for this. Importantly, waveforms during baseline vs licking periods had R^2^>0.9 correlation, also confirming that licking did not distort single unit waveforms. Bout and other lickometry measures were determined using custom-written programs in Python.

### Recording session methods

Before each experiment, an Omnetics headstage (Intan, CA) was connected to the implanted Multi Wire Array. The headstage was connected to a commutator (Doric Lenses Inc, Quebec, CA) via a standard serial peripheral interface (SPI) cable (Intan, CA). An additional SPI cable connected the commutator to the OpenEphys acquisition box (OpenEphys, MA) connected to a PC. Rats were then given ten minutes to acclimate before the experiment would begin.

Data was recorded at 30 kHz sampling rate using an Open Ephys (Cambridge, MA) system (acquisition board v2.2), run by Open Ephys software [66]. Since rats were well trained in the drinking paradigm, we recorded basal aINS firing for ~3-4 min before placing one alcohol bottle on the home cage. Following removal of the bottle, data was recorded for an additional two minutes.

All recordings were performed unilaterally in the right aINS since this region consistently activates in humans for intoxicant cues, drive for intake, and negative affects states [28,40].

### Extraction and identification of single unit waveforms

Raw, extracellular data was median referenced and subsequently spike-sorted using Kilosort 2 (KS2, with highpass filter at 600Hz). This was followed by exporting clustered units into Phy2 (https://github.com/cortex-lab/phy) for additional manual curation [67]. KS2 identified clusters were split into “good”, “mua”, or “noise.” “Good” clusters were further verified according to the following features: (1) distinct waveforms, (2) clear refractory period in crosscorrelograms, and (3) spikes forming a clear circular cluster grouping. For manual curation, clusters were split if they displayed two clear groups, and spikes without clear waveforms, or with no refractory period in cross-correlogram, were classified as noise and removed. Clusters labeled as “good” in Phy2 were then exported to custom-written MATLAB routines for further data analysis. Firing rates <0.33 Hz were not considered. We also calculate % inter-spike interval violations (spikes closer than 2ms) and exclude cells with >5% ISI violations. Putative single units were then analyzed using custom MATLAB scripts.

### Spike Train Analyses

Analyses use custom routines in MATLAB. Firing rate data was extracted from separate stages of the recording session in 500 ms bins. Basal firing was determined as the average across the 65 to 5 seconds before bottles were placed on the home cage, to 5 sec before rather than at the time of bottles on (because of possible anticipatory firing while watching the experimenter move in the room when getting the bottle). Basal firing in aINS cells could show strong variability across time, with firing interspersed with gaps in firing (see **Fig.2**). For this reason, standard deviation in the baseline was quite high, making standard Z-score methods impractical. Instead, we used a criterion using percent change for cells with >2Hz basal firing, and actual change for cells with <2Hz basal firing (where percent values were less meaningful). In particular, cells were considered to show increased firing in the given period if firing (initial licking or plateau licking) was 30% greater than basal (for >2Hz basal firing cells) or increased by 0.75Hz or more (for <2Hz basal firing cells). Similar criteria were used for cells showing decreased plateau firing, except with a 30% or 0.75Hz decrease relative to baseline. Other criterion showed similar patterns, although, with fewer cells at higher change criterion, there were trends (not shown).

We also examined the width of single unit waveforms, since in some regions GABAergic cortical neurons have thinner waveforms and higher basal firing (e.g. [61]). Overall, thinner waveforms were moderately but significantly correlated with higher baseline firing activity, but there were not clear delineations to indicate separate clusters of cells (**Suppl.Fig.1A,B**). There were also no differences across conditions in number of cells recorded, or waveform width or amplitude (**Suppl.Fig.1C-E**),

### Analyses and Statistics

Analyses were performed across firing neurons separate from rat identity: we have previously done this for alcohol-related behavioral microstructure [27,41], following other groups [68] including for analysis of aINS in vivo firing patterns [49]; Darevsky and Hopf (2020) has a detailed validation of such analyses.

Nearly all data were not normally distributed (determined with Shapiro-Wilk). Thus, statistical analyses were performed using the Kruskal-Wallis test (KW) with Dunn’s post-hoc, a non-parametric analog of one-way ANOVA, to compare across drinking conditions. To compare across two groups, we used Mann Whitney for unpaired and Wilcoxon for paired comparison was used as non-parametric t-test. Such analyses were performed using Graphpad Prism.

Principle Component Analysis (PCA) is commonly used as a dimensionality reduction tool that requires minimal assumptions of the data. Thus, we also determined firing patterns with PCA. An “omnibus” PCA utilized a matrix containing all data for all experimental conditions (drinking conditions), with population activity aligned at onset of drinking, and distributions of loadings of neurons on a given PC across experimental groups was examined with Kolmogorov-Smirnoff. Analyses used custom routines in R.

### Histology

Rats underwent paraformaldehyde (PFA) perfusion (details in Supplemental Material) to determine electrode placement. Prior to brain extraction, a stimulator (World Precision Instruments, FL) was used to run a small current through each electrode to lesion the location of the microwire endpoints. This led a small circular burn denoting the electrode tip (histology examples shown in **Suppl.Fig.2**).

While recording implants damaged overlying tissue, implants were only implanted unilaterally (in right aINS), and no behavioral changes were observed in locomotion or intake behavior due to implant. Similarly, previous studies using the same implant methods [58–61] did not observe non-specific behavioral effects of implantation.

## RESULTS

### Drinking Sessions Examined

Drinking sessions were 4 min long, with strong initial drinking (“load up”), and we examined several sessions of each drinking condition (AOD, AQ10, AQ60) in 15 rats in randomized order. To have most consistent comparison of in vivo firing patterns across drinking conditions, we selected two sessions of AOD, AQ10, and AQ60 from each rat, balancing high number of cells recorded in vivo and randomized order of exposure to drinking sessions. This yielded a total of 448 AOD, 378 AQ10, and 393 AQ60 aINS cells. Further, analyses were performed across firing neurons separate from rat identity: we have previously done this for alcohol-related behavioral microstructure [27,41], following other groups [68] including for analysis of aINS in vivo firing patterns [49]; Darevsky and Hopf (2020) has a detailed validation of such analyses.

### Drinking Behavior

Before addressing neural activity patterns, we examined behavioral responses which might indicate shifts in action strategy across AOD versus CLAD. Alcohol Drinking level (g/kg) was significantly reduced in AQ60 versus AOD or AQ10 (**Fig.1A**; KW *p*<0.0001; *p*<0.0001 AOD vs AQ60, *p*<0.05 AQ10 vs AQ60), consistent with higher challenge influencing ability to maintain alcohol intake. Difference across groups were also observed for total licks (**Fig.1B**, KW *p*=0.0194) and total licking time (**Fig.1C**, KW *p*=0.0234), with *p*<0.05 for AOD vs AQ60 for both measures. In contrast, there were no condition differences in bout number (**Fig.1D**, KW *p*=0.9131), average bout length (**Fig.1E**, KW *p*=0.1023), and length of pauses in licking (Inter-Bout Interval, IBI) (**Fig.1F**, KW *p*=0.4466). However, even with no average cross-condition differences, total licks and intake were highly correlated (**Fig.1G**, *p*<0.0001), and greater total licking did significantly correlate with average bout length for all three drinking conditions (**Fig.1H**, *ps*<0.003), with a similar trend for average bout length for g/kg intake but no relation between intake level and number of bouts or IBI length (**Suppl.Fig.3**). Finally, there were no condition differences in the rate of licking, shown for the first 60 licks (**Fig.1I**). Thus, there were limited behavioral differences across drinking conditions with the licking paradigm used here (home-cage licking from a bottle), although there were overall indicators of better organization of licking (e.g. longer average bouts) that did relate to higher drinking level. In addition, the similar IBI (length of pause during licking) and number of attempts to initiate licking (bout number) may suggest similar motivation to initiate alcohol consumption across drinking conditions, even if overall responding was somewhat less sustained under AQ60.

**Fig.1.**
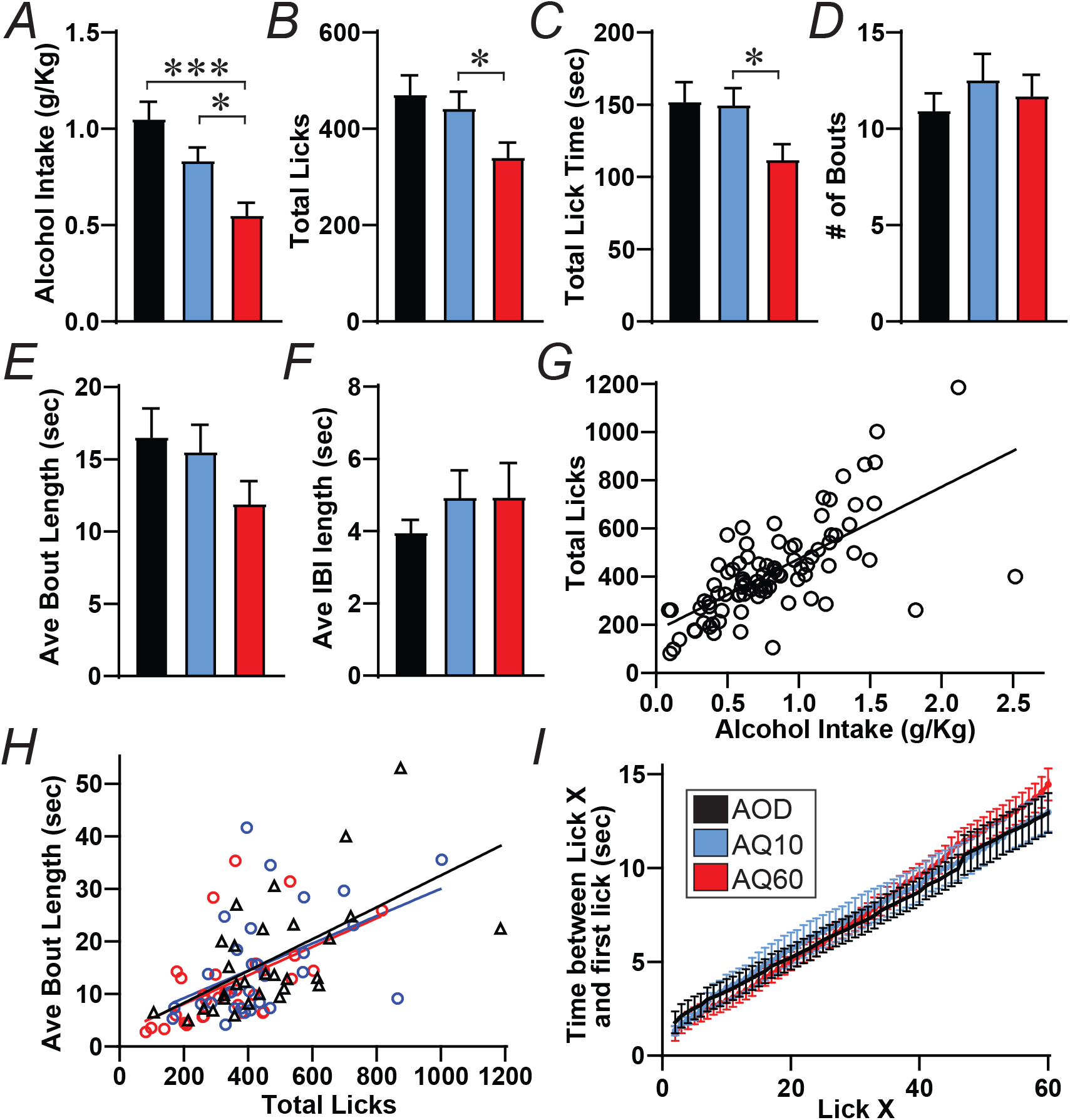
Behavioral patterns across AOD, AQ10, and AQ60 sessions. **(A-E)** averaged responding across drinking conditions. **(A)** Total intake levels (g/kg) were significantly reduced for AQ60 vs AOD (*p*<0.0001) and AQ60 vs AQ10 (*p*=0.0181) but not AQ10 vs AQ10 (*p*=0.3026). **(B)** Total licking was significantly reduced for AQ60 vs AQ10 (*p*=0.0242) but not AQ60 vs AOD (*p*=0.0989) or AQ10 vs AQ10 (*p*>0.9). **(C)** Total lick time was significantly reduced for AQ60 vs AQ10 (*p*=0.0499) but not AQ60 vs AOD (*p*=0.0563) or AQ10 vs AQ10 (*p*>0.9). Also no condition differences in **(D)** bout number, **(E)** ave bout length, or **(F)** length of Inter-Bout Interval (pause in licking). **(G)** Strong correlation between total intake and licking. **(H)** Total licks were significantly and positively correlated with average bout length for AOD (*p*=0.0005), AQ10 (*p*=0.0075) and AQ60 (*p*=0.0028). **(I)** No condition differences in the rate of licking, where y axis gives the time (in sec) between the first lick of the session and lick X across the session. *,*** *p*<0.05, *p*<0.001.

### Firing patterns of aINS cells with aspects of sustained responding during licking

Previous literature provides specific hypotheses regarding the importance of aINS activity for alcohol drinking, including (1) greater aINS activity for CLAD, and aINS activity corresponding with (2) initiation of responding and (3) sustained responding. Thus, we first examined firing patterns in cells sorted for a phenotype of strong firing increases at the onset of licking (Initial Response Phenotype, IRP, examples in **Fig.2A**). While IRP cells had no differences across conditions in basal, pre-lick firing (**Fig.3B**, KW *p*=0.4179) or firing during lick onset (average of first 2 sec, **Fig.3C**, KW *p*=0.2256), plateau firing averaged across the rest of the licking period was significantly greater under CLAD versus AOD. We defined the “macrobout” as the period including licking bouts and the brief pauses (IBIs), and firing in the first half of the IRP macrobout was significantly different across conditions (**Fig.3D**, KW *p*=0.0142), with both AQ10 and AQ60 plateau firing significantly greater than AOD (*p*<0.05). Significant differences across drinking conditions were also observed for the overall macrobout period (**Fig.3E**, KW *p*=0.0211, *p*<0.05 AOD vs AQ10) and the actual licking period (macrobout without IBIs) (**Fig.3F**, KW *p*=0.0188, *p*<0.05 AOD vs AQ60). We also compared firing by subtracting basal activity from firing during a given epoch for each cell. While lick-onset minus basal was not different across groups (**Fig.3G**, KW *p*=0.2827), significant differences were observed for first half licking minus basal (**Fig.3H**, KW *p*=0.0049, *p*<0.01 AOD vs AQ60) and actual lick period firing minus basal (**Fig.3I**, KW *p*=0.0064, *p*<0.01 AOD vs AQ60). Finally, firing post-licking quickly returned to basal levels and was similar across conditions (**Suppl.Fig.4A**, KW *p*=0.3952).Taken together, these data suggest that cells with strong firing increases at the onset of licking (IRP cells) showed significantly greater sustained firing during compulsion-like drinking conditions.

**Fig.2.**
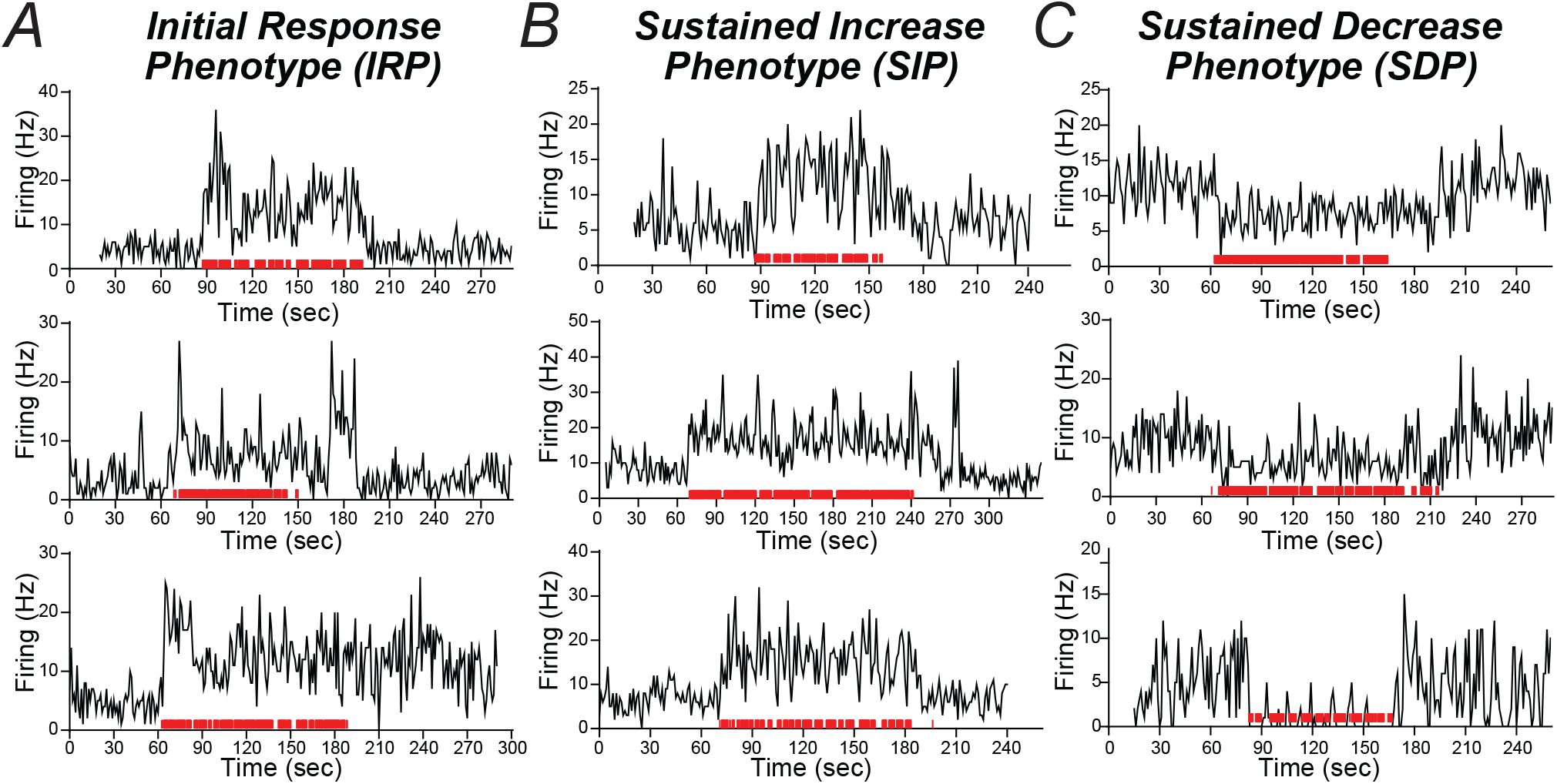
Examples of aINS cell firing, including **(A)** Initial Increase Phenotype (IRP), **(B)** Sustained Increase Phenotype (SIP), and **(C)** Sustained Decrease Phenotype (SDP). Red marks indicate licks, mostly fused together, e.g. with ~1000 licks in **(B)** top. For each phenotype, top, mid, and bottom examples are from AOD, AQ10, and AQ60.

**Fig.3.**
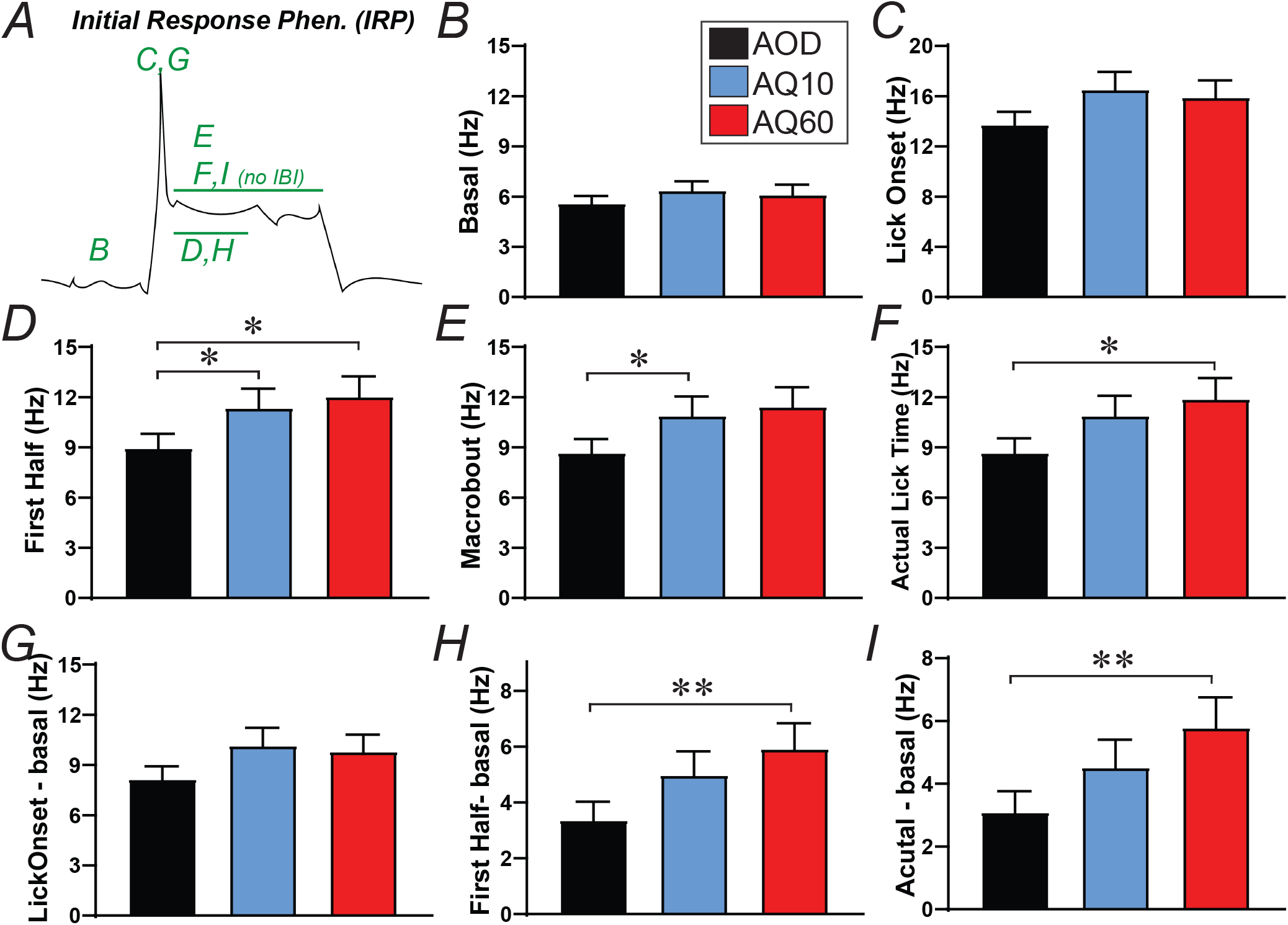
Initial Response Phenotype (IRP) cells had significantly greater plateau firing for CLAD conditions versus AOD. **(A)** Cartoon of IRP cell, with letters in green indicating the figure panel related to the particular letter. **(B)** Basal firing. **(C)** Lick onset firing. **(D)** Firing in first half of macrobout (the time period containing bouts and IBIs). Dunn’s posthoc *p*=0.0372 AOD vs AQ10, *p*=0.0453 AOD vs AQ60, *p*>0.9 AQ10 vs AQ60. **(E)** Firing in entire macrobout. Posthoc *p*=0.0391 AOD vs AQ10, *p*=0.0851 AOD vs AQ60, *p*>0.9 AQ10 vs AQ60. **(F)** Firing in actual licking (across bouts of licking). Post-hoc *p*=0.0668 AOD vs AQ10, *p*=0.0403 AOD vs AQ60, *p*>0.9 AQ10 vs AQ60. **(G)** Lick-onset minus basal firing. **(H)** First half minus basal firing. Posthoc *p*=0.0804 AOD vs AQ10, *p*=0.0056 AOD vs AQ60, *p*>0.9 AQ10 vs AQ60. **(I)** Actual lick period firing minus basal. Posthoc *p*=0.1804 AOD vs AQ10, *p*=0.0054 AOD vs AQ60, *p*>0.6 AQ10 vs AQ60. *,** *p*<0.05, *p*<0.01.

We next examined cells which showed increased firing across the licking period, which we call Sustained Increase Phenotype cells (SIP, examples in **Fig.2B**). Unlike cells aggregated for strong initial firing, there were no differences across conditions in SIP cells for any firing measure, including basal (**Fig.4B**, KW *p*=0.9632), lick onset (**Fig.4C**, KW *p*=0.9947), first half of the macrobout (**Fig.4D**, KW *p*=0.9199), whole macrobout (**Suppl.Fig.4B**, KW *p*=0.9330), actual licking period (**Fig.4E,** KW *p*=0.9678), or post-licking period (**Fig.4F**, KW *p*=0.7101). These results, along with **Fig.3**, lead to our speculation that it was not aINS plateau activity per se that contributed to sustaining compulsion-like drinking, but instead that it was plateau activity specifically in cells with strong initial firing (IRP cells) that sustained greater motivation for alcohol required under aversion-resistant, compulsion-like conditions (see Discussion).

**Fig.4.**
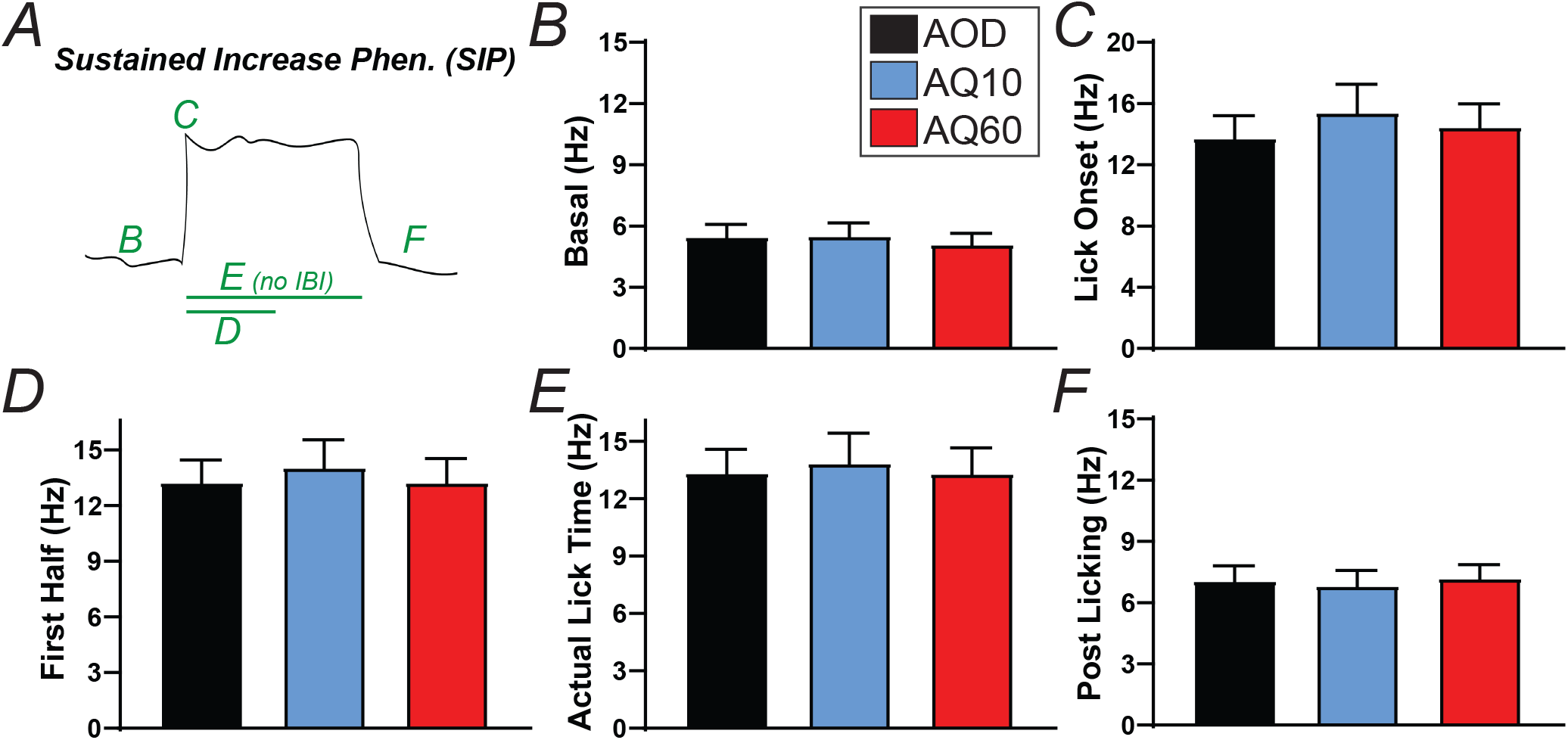
No condition differences in Sustained Increase Phenotype (SIP) **(A)** Cartoon of SIP cell, with letters in green indicating the figure panel related to the particular letter. **(B)** Basal firing. **(C)** Lick onset firing. **(D)** Firing in first half of macrobout. **(E)** Firing during actual licking. (F) Firing during post-licking period.

Both increases and decreases in firing have been described in aINS cells [48,49,69]. We did observe cells with decreased firing across the licking period, which we call Sustained Decrease Phenotype cells (SDP, examples in **Fig.2C**). However, there were no differences across drinking conditions in SDP cells in the firing level, including basal (**Fig.5B**, KW *p*=0.7597), lick onset (**Fig.5C**, KW *p*=0.2313), actual lick period (**Fig.5D**, KW *p*=0.6311), changes in firing between actual lick period and basal (**Fig.5E**, KW *p*=0.7179), post-licking (**Fig.5F**, KW *p*=0.8682), or other measures (**Suppl.Fig.4C,D**). Thus, while SDP cells might contribute to drinking level (below), SDP activity likely did not differentially contribute to CLAD versus AOD.

**Fig.5.**
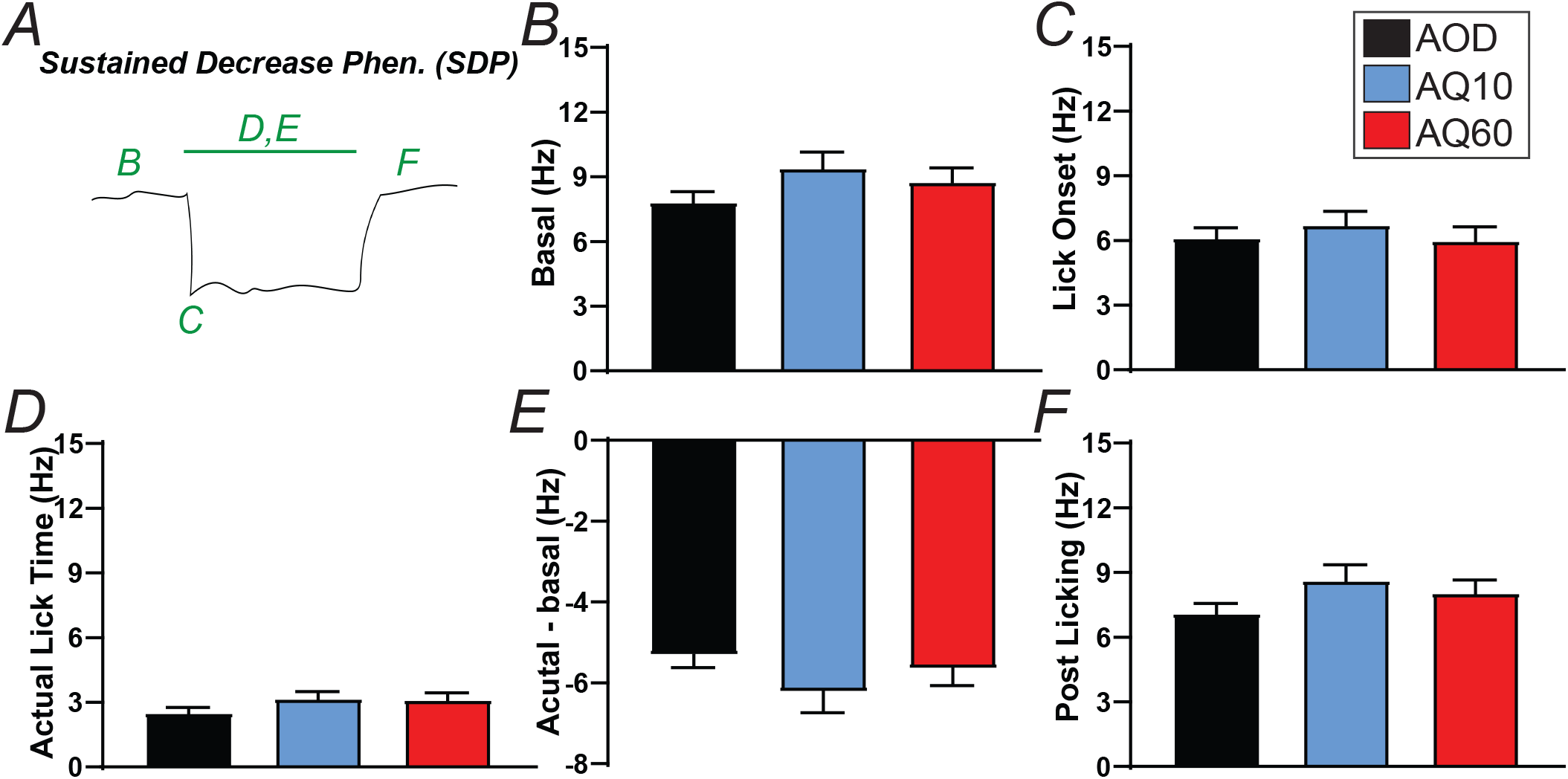
No condition differences in Sustained Decrease Phenotype (SDP) **(A)** Cartoon of SDP cell, with letters in green indicating the figure panel related to the particular letter. **(B)** Basal firing. **(C)** Lick onset firing. **(D)** Firing during actual licking. **(E)** Firing during actual licking period minus basal. **(F)** Firing during post-licking period.

Other studies have found shifts in the number of cells with increased or decreased firing across conditions [49,69]. However, we found no differences across drinking conditions in the incidence of different cell phenotypes, with ~35% IRP, ~25% SIP, and ~40% SDP cells across conditions (**Suppl.Fig.5**). However, we note that IRPs were defined by increased firing at lick onset, and thus some IRP also exhibited sustained increases or decreases in firing (e.g. the elevated sustained firing that differed across drinking conditions in **Fig.3**). Thus, we also examined overlap of SIP and SDP within IRP cells. There were more IRP-SIP (~50-60% of IRP cells) than IRP-SDP (~15-20% of IRP cells), with no condition differences (**Suppl.Fig.5**). Thus, shifts in aINS that could sustain CLAD did not involve overall changes in incidence of the three cell phenotypes.

Together, these findings suggest that there were limited differences in sustained firing patterns across drinking conditions, and thus that particular, more nuanced aspects of firing differences could promote aversion-resistant drinking (see Discussion). To better understand overall firing changes, we performed a Principal Component Analysis of session-long activity patterns across all cells. PC1 explained ~30% of variation in firing, and was a sustained change across the licking period which did not diff among the drinking conditions (**Suppl.Fig.6**, Kolmogorov-Smirnov *ps*>0.8). Other PCs explained <5% or less of variability (PC2 and PC3 are shown in **Suppl.Fig.6**). Thus, these unbiased assessments indicate that any aINS firing changes that could promote CLAD vs AOD did not reflect larger, macroscopic changes in aINS activation. Instead, as described below, we speculate that CLAD involves a more nuanced shift in cognitive/emotional shift in action strategy in face of challenge, which works in parallel with a separable intake-promoting activity (following section).

### Measures related to total responding for alcohol

We next examined whether any aspects of firing patterns predicted the level of alcohol responding. Interestingly, the only aspect of firing that predicted total licking was the percent of cells with SDP phenotype (**Fig.6A** *p*=0.0027 collapsed across drinking conditions). When separated by drinking condition (**Fig.6B**), %SDP cells in a session was positively correlated with total licks for AOD (*p*=0.0271), and a trend for AQ60 (*p=*0.0519), although not for AQ10 (*p*=0.4573) (see **Suppl.Fig.7** for g/kg versus cell type incidence). However, total licks in a session did not correlate with % IRP cells (**Fig.6C**, *ps*>0.19) or %SIP cells (**Fig.6D**, *ps*>0.4). Thus, only SDP cells shown a relationship to total intake.

**Fig.6.**
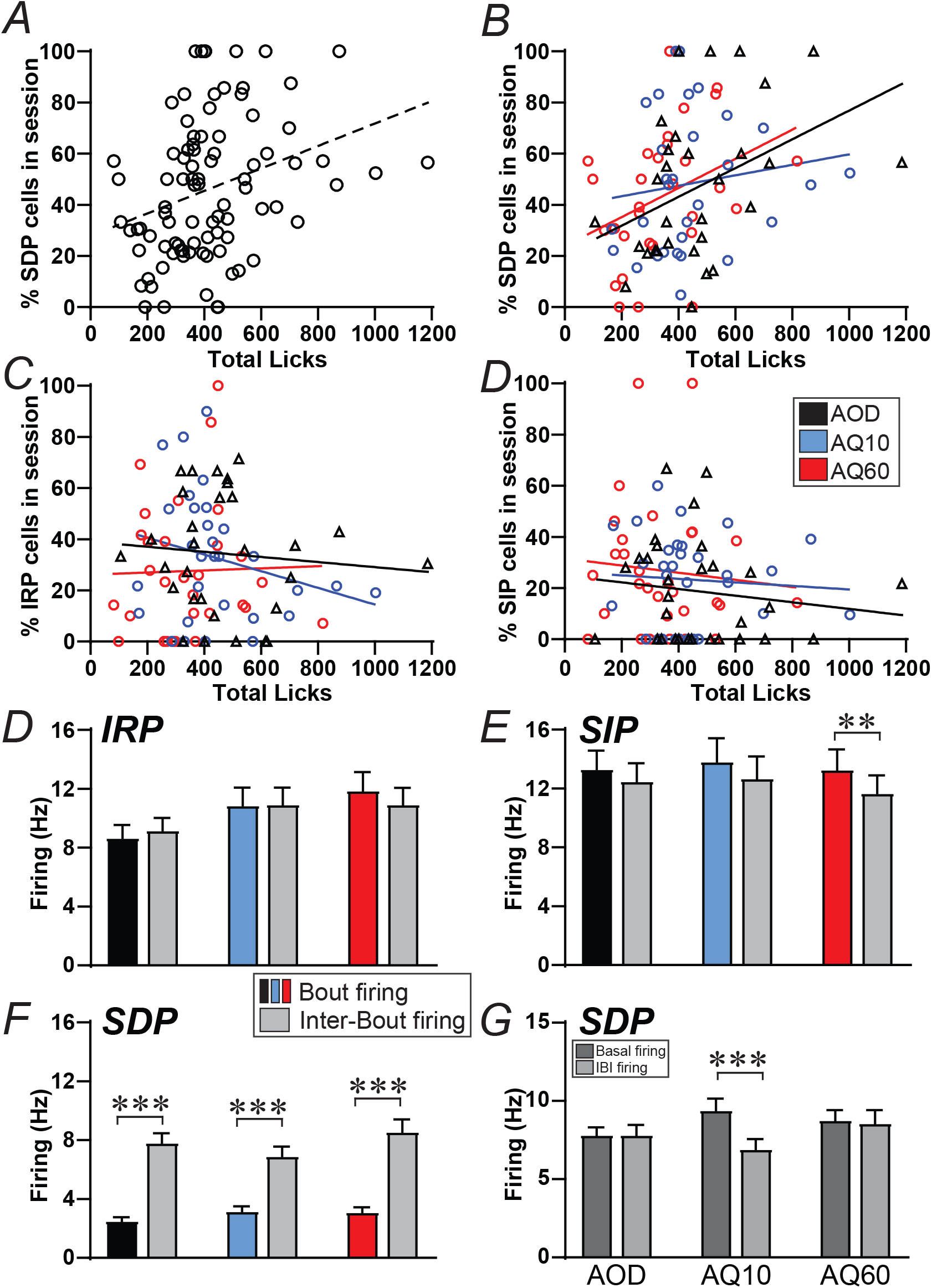
SDP cells: abundance predicts intake level, although SDP cells to not maintain reduced firing during pauses in licking. **(A,B)** Total licking is correlated with greater %SDP cells in a session for **(A)** all conditions collapsed, and **(B)** for AOD and AQ60 alone. **(C,D)** No relation between total licking and **(C)** %IRP cells or **(D)** %SIP cells within a session. **(E,F)** IRP **(E)** and SIP **(F)** cells maintain their plateau responding during pauses in licking (IBIs). **(G)** SDP cell firing significantly increases during pauses in licking in all drinking conditions. **(H)** SDP cell IBI firing is not different from basal for AOD and AQ10. **,*** *p*<0.01, *p*<0.001.

Another interesting aspect of SDP cells related to the firing level during pauses in licking, the Inter-Bout Intervals (IBIs). IRP cells maintained their plateau firing level during IBIs (**Fig.6E**), such that average IBI firing did not differ from actual lick period firing for any drinking conditions (Wilcoxon AOD *p*=0.0957, AQ10 *p*=0.5027, AQ60 *p*=0.0776). SIP cells showed no difference in IBI and actual lick period firing for AOD (*p*=0.1098) and (AQ10 *p*=0.1488), with a slight yet significant difference for AQ60 (*p*=0.0023) (**Fig.6F**). In contrast, IBI and actual lick period firing were strongly different for SDP cells from all drinking conditions (**Fig.6G**, *ps*<0.0001), and IBI firing level was not different from basal (pre-lick firing) for AOD (*p*=0.1455) or AQ60 (*p*= 0.4961), although it was different for AQ10 (*p*=0.0009) (**Fig.6H**). There were also no differences in IBI activity across conditions for any cell phenotype (KW; IRP *p*=0.2089; SIP *p*=0.8764 SDP *p*=0.1992). Thus, drinking level overall was only related to the number of SDP cells within a session, while only IRP and SIP cells maintained responding across pauses in licking. As described below, we interpret these findings within a hypothesis where IRP cells (which showed session-level changes in CLAD-related plateau firing) represented the overall action strategy to maintain aversion-resistant responding, assuring that sufficient intake occurred under CLAD conditions. In contrast, SDP cells primarily reflected actual licking, and thus we speculate they contributed to intake level overall.

## DISCUSSION

Compulsive-like alcohol drinking (CLAD), where intake persists despite adverse consequences, can be a core facet of Alcohol Use Disorder and major obstacle to treatment. Further, despite advancements of our understanding of mechanisms underlying CLAD (e.g. [34,42,43,70]), much remains unknown. Previously, we described that aINS, a central mediator of emotion, motivation, and interoception, promotes CLAD in rodents, and heavy human drinkers show similar insula-circuit recruitment under compulsion-like conditions. However, global aINS inhibition reduces CLAD and alcohol-only drinking (AOD), perhaps suggesting parallel aINS pathways for CLAD versus AOD. One major obstacle to understanding aINS importance is the lack of information on in vivo aINS firing patterns, which could reveal critical insights into the nature of aINS activity that promotes alcohol intake. Here, we used 32-electrode multi-wire electrophysiology to record single-unit activity in right aINS from 15 rats during brief sessions of AOD or CLAD (10mg/L or 60mg/L quinine in alcohol). aINS is implicated in both initiating and sustaining motivated responding, and we find a confluence of these patterns. Neurons with a phenotype of sustained firing increases (SIP) or decreases (SDP) showed no firing differences across drinking conditions. In contrast, aINS neurons with a phenotype of strong firing increase at initiation of intake (IRP) showed significantly greater activity across the rest of licking during CLAD relative to AOD. This concurs with our previous findings that rats quickly assess that they are drinking under CLAD conditions and adjust their response patterns to a more aversion-resistant action strategy (below). There were also no condition-related differences in abundance in these firing phenotypes. Further, total responding in a given session correlated with abundance of SDP cells, but not IRP or SIP. In addition, SDP firing returned to baseline during pauses in licking, while IRP and SIP had sustained responding through pauses in licking, suggesting that SDP cells contributed to actual licking, while IRP and SIP more likely reflected a sustained action plan. These comport with our previous model [27,41] that CLAD represents a singular, session-long strategy to assure that sufficient intake occurs, but more detached from moment-to-moment responding (see below), which we speculate would be mediated by IRP. Together, INS cells with a particular strong firing at licking onset show greater sustained responding under compulsion-like conditions, while other cells likely promote drinking more generally, providing important new information about how aINS activity could sustain different aspects of alcohol consumption.

We focus here on aINS since it is a central mediator of emotion, motivation, and interoception, and is broadly implicated in expression of addiction and other neuropsychiatric conditions [28]. The aINS, through regulation of the salience network, mediates rapid identification and responding to important situations, and regulates accompanying emotional states [28–32]. Further, human aINS activity is linked to many aspects of alcohol drinking [28], including where alcohol cue- [71,72] and negative affect-induced [73] aINS activation predict real-world drinking (see also [74]), and aINS activation to alcohol images predicts later transition to heavy drinking [75]. Also, aINS changes are linked to worse behavioral control in AUD [76,77], while therapies that reduce drinking decrease aINS activity and/or connectivity (e.g. [78,79]). Thus, aINS likely mediates factors that promote pathological drinking.

We also focus on aINS because of critical preclinical work indicating the region’s importance for many addiction-related behaviors. We found that aINS-NAcb core in rat is critical for compulsion-like responding for alcohol, using quinine or footshock as the aversive consequence [33] (see also aINS-NAcb for compulsion-like responding for sugar [80]), and that aINS-Locus Coeruleus area is also critical for rat CLAD [34], while neither pathway is necessary to regulate AOD. Further, modulating putative aINS GABA signaling significantly decreases CLAD but not AOD in rat [34] and mouse [35]. In concurrence, aINS circuit activity in heavy drinking humans is related both to the level of punishment-resistant responding for alcohol and subjective compulsivity for alcohol [38]. Further, in women with AUD, imaging high-risk drinking contexts activate aINS more than low-risk intake situations [37]. Importantly, recent studies have also helped establish the relation between human and rodent aINS [28,81,82], overall supporting the speculation that at least some aspects of aINS signaling have a more specific relation to conflict-resistant responding (see Suppl. Discussion). In contrast, we find that global inhibition of aINS, targeting cell activity more broadly or alpha1-noradrenergic receptors specifically, suppresses both AOD and CLAD [34]. Similarly, aINS activity in humans can relate to alcohol intake more generally (above), and global aINS inhibition in rodents reduces self-administration of several other substances, including nicotine [83], cocaine ([84,85] but see [86]), and palatable food [87], while inhibiting the adjacent lateral OFC has no effect on CLAD [34,88]. Together, these findings support the possibility that different aINS mechanisms promote AOD versus CLAD, along with behavioral evidence that punishment-resistance for alcohol does not correlate with AOD level [21,36], as seen for other drugs (e.g. [89,90]). Further, while others observed that aINS-NAcb does not regulate AOD [91], Jaramillo and colleagues find that aINS-NAcb inhibition reduces operant self-administration of sweetened alcohol and regulates alcohol interoception [39,92,93]. Also, global aINS inhibition does not impact alcohol relapse in early withdrawal, although does mediate punishment-resistant alcohol seeking [36]. Some studies use shorter overall alcohol intake history relative to our work, and the long-term intake we utilize may impact mechanisms of regular, alcohol-only drinking. Thus, additional work is needed to address whether different procedures and/or other factors might contribute to potentially divergent findings. Nonetheless, inhibiting aINS or aINS projections does not impact sweet fluid [33,34,93,94] or water intake [87,91] (but see [95]), simpler operant learning and responding [83,96,97], or locomotion [34,36,87,91,93,98,99], suggesting important selectivity in aINS contributions to motivated responding (although activating aINS does reduce consumption of several substances [91,95,100,101]).

Since different aINS cell firing phenotypes may relate to varied aspects of alcohol drinking, with IRP plateau activity related to maintaining aversion-resistant responding, and the incidence of SDP cells related to greater intake level overall, one key question is the potential nature of these two aINS cell types. Indeed, while ~40% of cells were SDP, <20% of IRP cells were also SDP, suggesting at least partial separation of the firing phenotypes. One simpler possibility is that aINS cells with different projection targets reflect the different firing phenotypes. For example, in many cortical areas, pyramidal neurons fall into three general classes, pyramidal tract (primarily projections to subcortical areas), intratelencephalic (primarily to cortical areas), and thalamic projectors [102]. In this regard, in rat there is little overlap between aINS cells that project to subcortical regions vs cortical areas [103], while one study estimates co-projection of aINS cells to both amygdala and NAcb core at <10% [104] (see also [101]). Thus, different aINS projections may come from relatively distinct subgroup of aINS cells, with potential for different firing patterns. As noted above, aINS-NAcb has been related to several CLAD behaviors, and a recent preprint implicates it in expression of chronic pain [105]. Additionally, we find that that aINS-Locus Coeruleus Area contributes more to CLAD than AOD [34], similar to certain aspects of NMDA (see [23]) and orexin (see [106]) receptor signaling. However, interpreting such findings, especially the particular behavioral importance of a given aINS projection, requires some caution. For example, aINS-BLA contributes to appetitive drives [107–109], but also in fear learning [110], taste aversion [111,112], aversive visceral stimuli [95], and glutamate receptor adaptations that contribute to aversive symptoms with alcohol withdrawal [113]. Indeed, the Salience Network is at core about readiness to respond under high-importance situations, and thus can contribute to both negative and positive conditions (see [28,40]). In addition, some aINS connections have received very little attention, e.g. even though aINS-cortical outputs can be quite large [28,81,82,114,115], we are only aware of a single publication directly examining the behavioral importance of aINS outputs to other cortical areas, where aINS-dmPFC (also aINS-BLA and dmPFC-aINS) are critical for expression of avoidance of novel tastes, while learning to habituate only requires aINS-dmPFC [116]. Thus, the identity of aINS cell types with different in vivo firing patterns remains an area of critical and unmet interest.

Identifying actual neuronal firing patterns is essential for understanding how aINS may differentially contribute to CLAD vs AOD (and other behaviors). Together, our main findings suggest that IRP activity could sustain CLAD more than AOD, and that SDP cells are related to greater intake overall. Importantly, we interpret these findings within previous work indicating aINS importance for both initiating and sustaining behavior, and where CLAD likely involves a cognitive/emotional strategy which allows persistent responding under challenge while reducing the impact of adverse contingencies [27,41]. Indeed, while aINS and Salience Network are implicated in initiating many motivated behaviors (reviewed in [28]), aINS is also likely important for sustaining motivated responding. Persisting aINS signals reflect the main goals of a task [44,45] and persisting emotion [46] in humans, and the long-term reward value of a context in non-human primates [47]. Although there are quite limited studies in rodents, Guillem and colleagues (2010) [48] show hours-long plateau firing across a 3hr cocaine intake session in half of aINS cells (see also [49]), and aINS maintains cue responding across longer delays [117]. Thus, aINS circuits can support both initiation and sustaining of motivated responding.

In particular, we have described a “Head Down and Push Model” of CLAD, based in part on behavioral evidence that rats quickly determine whether they are drinking alcohol-only or alcohol with aversion, and adjust responding accordingly [27,41].. This model is akin to “intake defense” (e.g. [50]), where the goal is to assure that sufficient intake occurs rather than specifically titrating intake level to motivation level under conditions of challenge (such as water deprivation or hunger, see also Suppl. Discussions in [27,41]). In this model, one would predict neuronal activity related to CLAD strategy across the session, which would be critical for CLAD responding per se but more unlinked from the exact intake level (as long as “enough” intake occurred). It is also interesting, in this model, that the incidence of SDP cells in a given session correlated with higher intake, and yet SDP cells did not sustain plateau-related firing during pauses in licking, unlike IRP and SIP cells which did maintain firing across such pauses. Thus, a priori, IRP and SIP were more likely candidates for encoding a main CLAD action strategy, with SDP more related to actual licking activity. In this regard, binge-prone rats (more prone to palatable food binging) have greater activity in several regions during pauses in licking, when compared to binge-resistant rats, which the authors interpret as maintained expectation for reward in food bingers [54]. Also, in our previous studies, the timing of the first lick of a session did not differ across drinking conditions, but, after the first lick occurred, responding happened significantly sooner under CLAD versus AOD; in addition, under moderate challenge, many aspects of lick responding became less variable, and while some aspects of reduced variability were retained under higher challenge, others were lost in concert with reduced intake [27,41]. However, these previous studies used different licking requirements, with a more complicated geometry, than utilized here (licking in a Med Associated operant box, where the rat had to rear slightly and insert it’s nose into a space to lick, compared to simple home-cage bottle drinking in the present study). Nonetheless, our present and previous studies together support the concept of a singular, session-long CLAD action strategy to sustain responding to persist in the face of challenge while minimizing attention to and impact of negative consequences (see [27,41], and [118] for a related conceptualization of maintaining “adaptive response to stress” which involves Locus Coeruleus signaling).

One challenge to our interpretation of IRP linking initial evaluation of response conditions to sustained shifts in action strategy is that activity level at lick onset was not different across drinking conditions. One possibility is that aINS was important for an initial attentional response (with aINS strongly linked to attentional processing [28,40]), while another brain area (such as NAcb or Locus Coeruleus) determined whether the drinking condition required additional attention across further responding, which would be relayed to the insula to produce greater sustained firing to maintain CLAD. Alternately, we note that there was a trend for greater lick-onset firing under CLAD conditions, and further studies might identify that specific aINS projections with IRP firing patterns also have greater lick-onset activity under CLAD conditions.

The relation of SDP cell number to consumption that we observed here might relate to similar to consumption-related firing decreases observed in other regions (e.g. [51,52]) but not all (e.g. [53,54]). Such studies vary in a number of factors, including food deprivation and the nature of the food reward (sometimes highly palatable). In addition, it may seem counterintuitive that having more decrease firing cells relates to greater consumption level, while more global aINS inhibition reduces intake. Since different aINS projections can have different and potentially divergent impacts on behavior, we speculate that future work that uncovers the nature of aINS SDP cells will shed additional light on how these cells might promote alcohol consumption.

One more general issue is whether rodent CLAD models involve mechanisms related to human problem intake. For example, rodent CLAD paradigms have been challenged because the negative consequences in humans often occur in the future, which is unlike the acute adversity in rodent models. However, we favor a different perspective, where treatment-seeking humans experience the negative consequences more acutely [16] (they are “actively struggling with the balance between known bad outcomes and the desire to drink” [119]), and we have always considered our findings more relevant for treatment seekers. Indeed, our findings of aINS circuit importance for CLAD but not AOD comports with clinical theories that compulsion-like responding in human addiction is at core about conflict (between desires for intoxicant and avoiding negative consequences), and that it is the presence of this conflict that recruits cortical conflict-processing systems including the aINS and Salience Network [19,120]. Thus, putative human treatments based on our rodent findings would only be of utility for humans actively struggling with their conflict over intake. For these reasons and others, some clinicians have expressed support for the value of rodent CLAD models [20,38,42,43,120,121]. This is also supported by cross-species importance of aINS signaling [33,35-38,55] and perhaps NMDA receptors (see [23]) for compulsion-like responding for alcohol, and where people with AUD often continue to drink even as negative consequences increase [16–19]. Nonetheless, one group found that people with AUD were not more insensitive to cost [122,123] which could question the importance of consequence-resistance. However, intake despite negative consequences features prominently in the DSM-V definition of AUD (e.g. [14,26]), and other groups do find less sensitivity to cost with AUD [124–126], while greater drinking is associated with more alcohol problems [11–14]. Thus, some authors [20,121] have emphasized that, from a clinician’s perspective, multiple factors can contribute to AUD expression [127–133], including compulsion-like drives. In addition, the intermittent access alcohol-drinking model we utilize exhibits several features which could be considered related to human AUD, including escalation of intake [25,57], sensitivity to compounds that reduce human drinking [57], withdrawal symptoms (although moderate) [134,135], and front loading (indicating motivation for alcohol [27,41,136,137]), in addition to compulsion-like responding.

Another major limitation of our study is that we did not examine aINS firing patterns in female drinkers. Problem alcohol drinking in women has risen dramatically in recent years [1,138-142], and women can have greater alcohol problems including negative impacts on health [143–146] and may develop dependence faster [147–149]. Also, while female subjects have historically been excluded from many studies, a number of studies find important sex differences in brain mechanisms (reviewed in [119,145,150-153]) which are critical to take into consideration when developing personalized treatments [119,154,155]. It is interesting that the limited extant human studies link aINS circuitry to compulsion for alcohol in both sexes [37,38], although considerable future work will be required to better understand sex-different and -similar brain mechanisms for addiction and other cognitive/emotional conditions.

Problem alcohol consumption is associated with major social, physical, and financial harms, and, with the considerable evidence linking aINS to expression of addiction-related states, it is critical to understand how aINS firing patterns in vivo might contribute to alcohol-only and compulsion-like alcohol consumption. Here, by examining firing patterns across a larger sample of subjects, we discovered firing patterns that concur with previous hypotheses about how aINS could promote different aspects of alcohol responding under compulsion-like vs alcohol-only conditions. Specifically, CLAD would involve aINS cells with sustained encoding across the alcohol-intake session, which we speculate reflected a session-long strategy to sustain aversion-resistant intake. It is particularly interesting that this firing phenotype (IRP) had strong firing at lick onset, since we propose that rats quickly evaluate the licking conditions (alcohol-only or aversion-related) and rapidly adjust action strategy. In contrast, cells with decreased firing (SDP) did not maintain sustained firing during pauses in licking, but the abundance of such cells did predict drinking level overall. Thus, we provide valuable, novel information about aINS activity patterns that could promote different aspects of pathological alcohol consumption.

## AUTHOR CONTRIBUTIONS

PS and FWH designed the experiments; PS, MM, SK, RF, TDOS, and SW helped perform the experiments; PS, MM, CL and FWH analyzed the data; PS and FWH wrote a first draft of the manuscript, all authors edited the manuscript and approved the final version.

## Supplemental Materials

### Supplemental Methods

Rats were anesthetized before transcardial perfusion with phosphate buffer saline (PBS, 250 ml) then 4% paraformaldehyde (PFA) in PBS. Brains were extracted and post-fixed overnight in 4% PFA. Brains were reconstituted in 30% sucrose in PBS for 3 days before being sectioned using a cryostat at a thickness of 40 um. Brain slices were mounted onto positively charged frosted slides and dried overnight. The slices were then stained using a cresyl violet (Sigma-Aldrich, MO) and imaged at 2.5x. Placement of the microwires was determined by comparison of cresyl violet stained tissue against a Rat Brain Atlas (Paxinos & Watson, 1997).

### Supplemental Discussion

Consistent with aINS importance for CLAD, many human studies find greater aINS with higher attentional and response demand [1–6], which is related behaviorally to less interference from conflict [7,8] and intrusive thoughts [9,10], better behavioral control under emotion [11,12], and pro-risk responding [13–16]. However, aINS activity is also seen with simpler and positive events [17,18]; in this regard, mouse studies find aINS cFos activity for both AOD and CLAD, but with greater cFos for CLAD [19].

We also note that, while we hypothesize that aINS behavioral importance will be accompanied by parallel aINS firing changes, these may not always be linked. For example, in [20], aINS firing for task events does not change during aversion conditioning, even though aINS mediates basic conditioned aversion [21,22]. On the other hand, aINS neurons fire for different tastants [23–26], but aINS inhibition does not impair many aspects of taste discrimination [27,28]. Thus, aINS could promote behavior even without differential aINS firing.

### Suppl. Fig. Legends

**Suppl.Fig.1.**
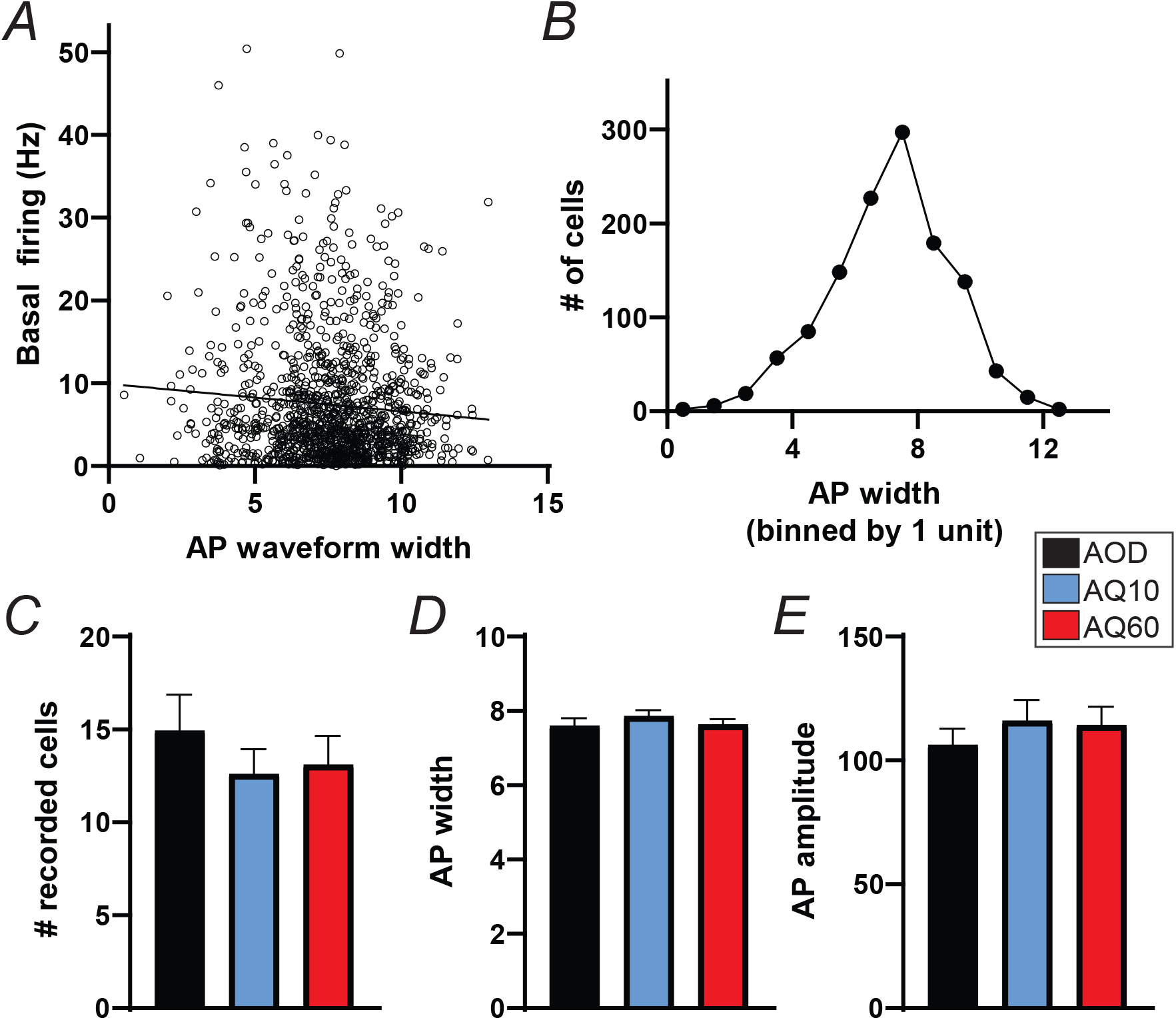
In vivo waveform parameters. In some cortical regions, GABAergic interneurons can be distinguished by narrower waveform. **(A)** Overall, thinner waveforms were moderately but significantly correlated with higher baseline firing activity (**Suppl.Fig.1A**, *p*=0.0038, R^2^=0.0069), but **(B)** there were not clear delineations to indicate separate clusters of cells. This is consistent with cells with thinner waveforms having higher firing rates, although the effects are modest and further studies would be required to characterize potential firing patterns of aINS interneurons. There were also no differences across conditions in the average per session of **(C)** number of cells recorded (KW *p*=0.8242), **(D)** waveform width (KW *p*=0.5684), or **(E)** waveform amplitude (KW *p*=0.7288).

**Suppl.Fig.2.**
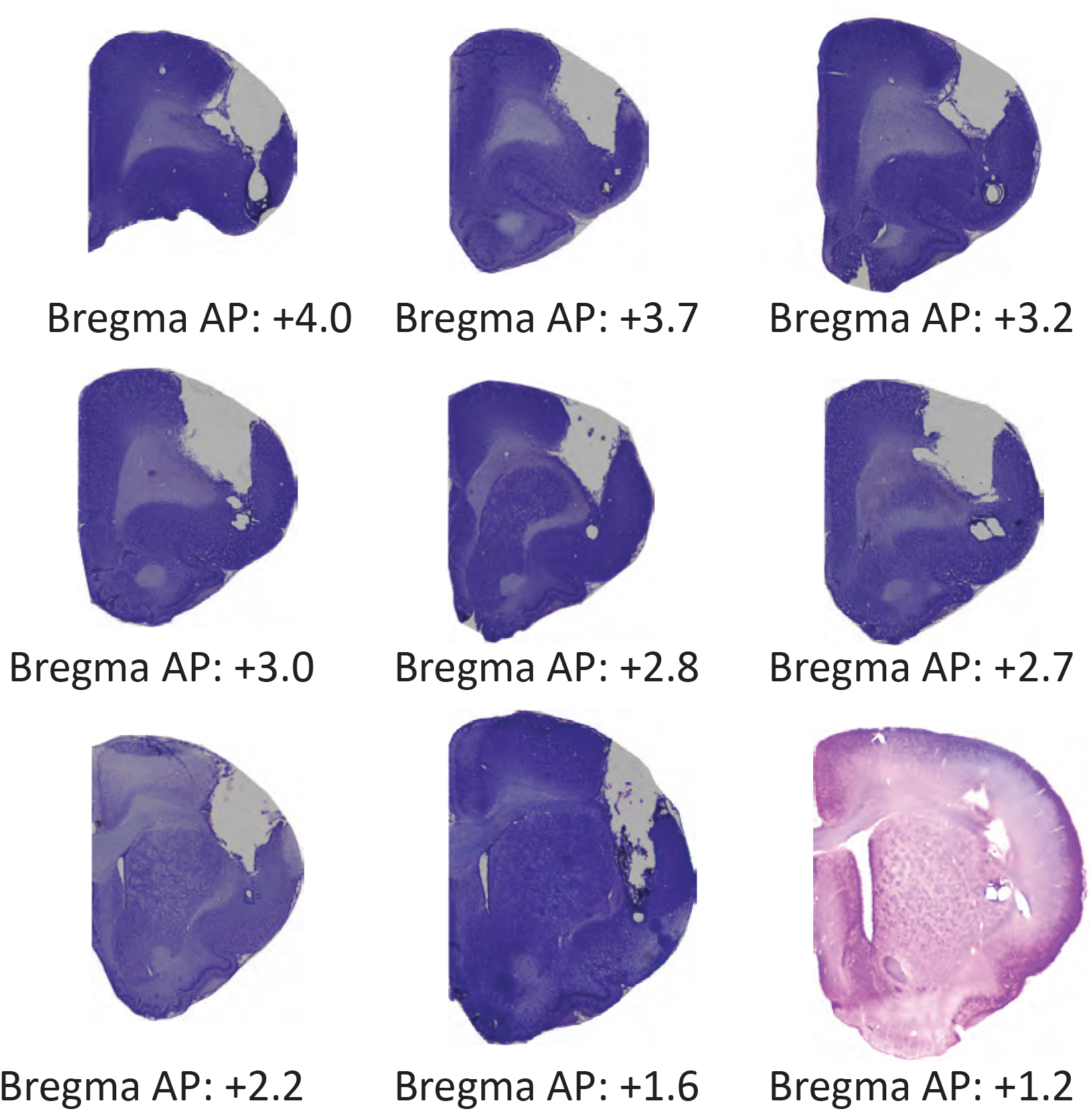
Examples of recording electrode placements in aINS. Nissl stains. At the end of each study, after PFA fixation, a small current is run through each electrode, with the goal of creating a small circular burn (arrows) noting the electrode tip.

**Suppl.Fig.3.**
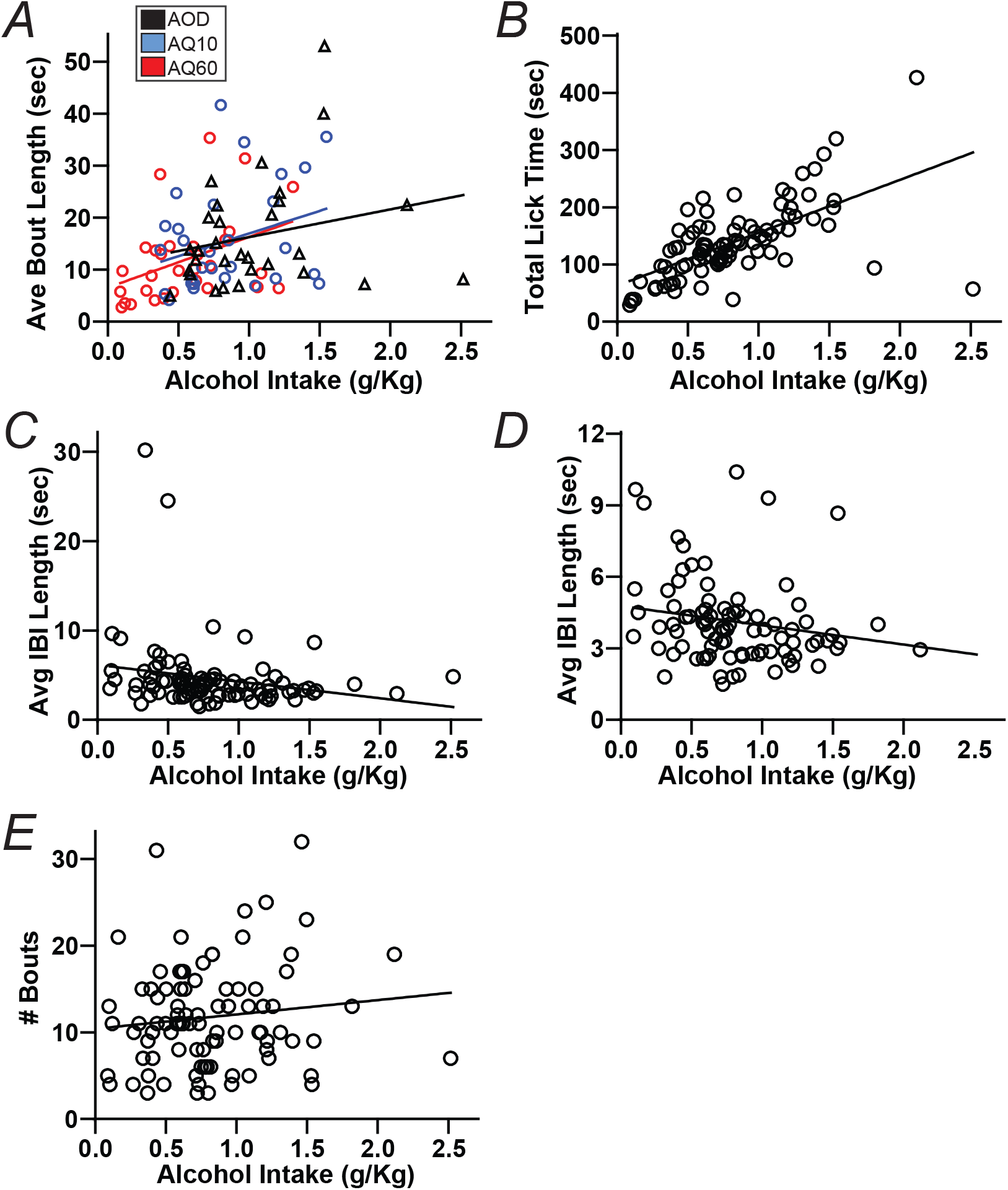
Session Level Correlations. **(A)** Total intake level significantly correlated with ave bout length for AQ60 (*p*=0.0304) but not AOD (*p*=0.2042) or AQ10 (*p*=0.0987). **(B)** Intake level strongly correlated with total time of licking (*p*<0.0001). **(C,D)** Intake level did correlate ave IBI length with all data points **(C)** (*p*=0.0442) but no longer did so after removing two high outliers **(D)** (*p*=0.0627). **(E)** Intake level did not correlate with bout number (*p*=0.2518). For **(BE)** session measures are collapsed across drinking conditions. See also **Suppl. Table 1** for relations between alcohol intake level and a given session-level measure, collapsed across all sessions.

**Suppl.Fig.4. Some firing relations across conditions. (A)** Post-lick firing in IRP cells. **(B)** Macrobout firing in SIP cells. (**C**) Macrobout firing in SDP cells (KW *p*=0.8865). **(D)** First half of macrobout firing in SDP cells (KW *p*=0.8135).

**Suppl.Fig.5.**
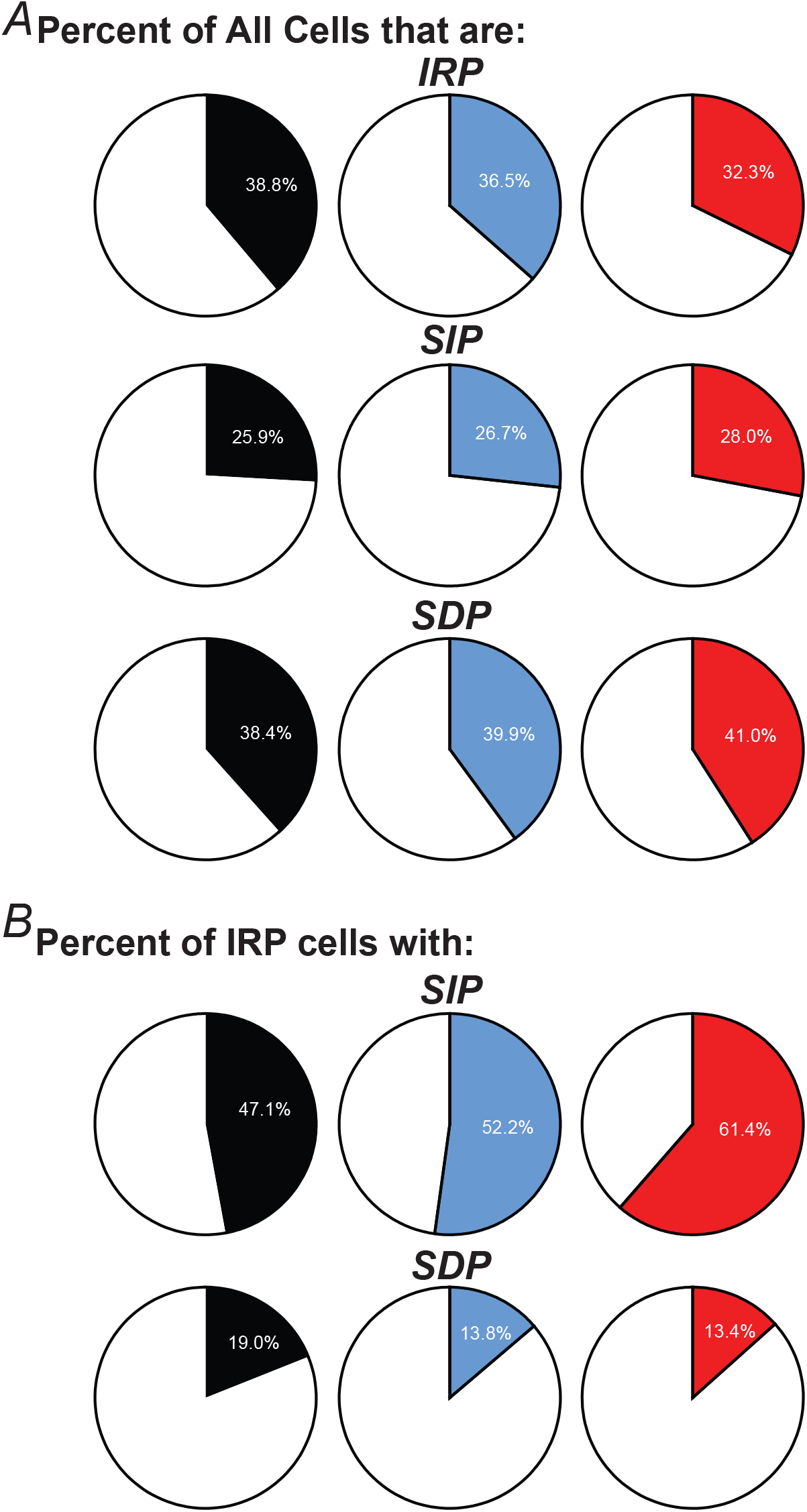
Incidence of different cell phenotypes, relative to all neurons recorded in a given session. **(A)** % of cells in a session with IRP, SIP, or SDP. **(B)** % of IRB cells in a session with SIP or SDP.

**Suppl.Fig.6.**
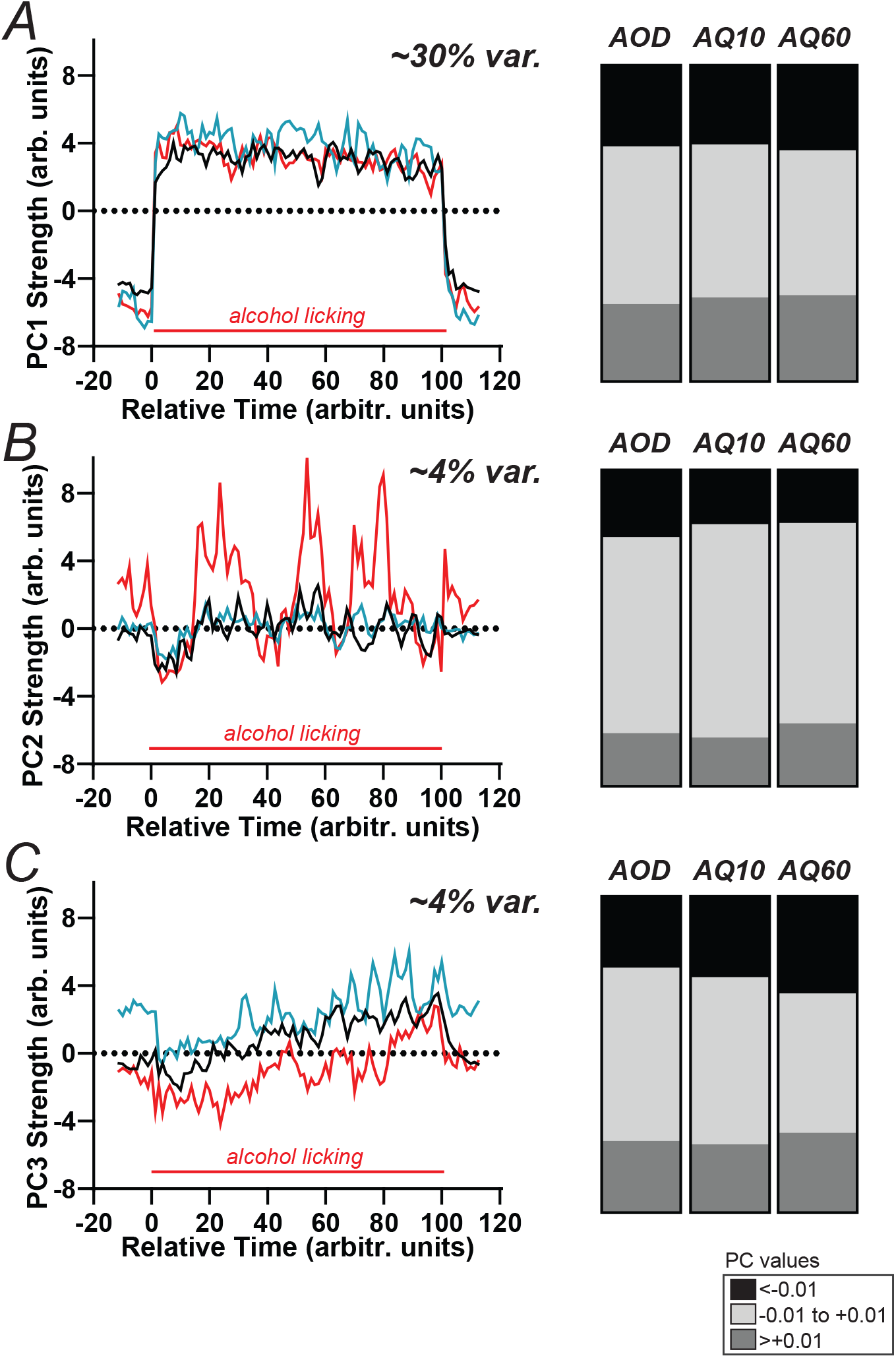
PCA of overall firing patterns across the licking period, compared between drinking conditions. In order to compare across sessions, which had different durations of consumption, we first normalized the time axis for each session so that firing was assessed across an equal number of intervals (100) for each session. This allowed comparison of firing changes across sessions in a common “session-passing” frame. We examined how drinking condition might impact contributions to each PC with Kolmogorov-Smirnov test to examine differential pattern of loading on cells. Left column examines the 3 top PCs across the entire licking session. For PC1 (~29.7% of variance) there were no condition differences: Alc vs AQ10: D(825)=0.0304, *p*=0.9904; Alc vs AQ60: D(840)=0.0426, *p*=0.8344; AQ10 vs AQ60: D(771)=0.0375, *p*=0.9453. For PC2 (~4.4% of variance) there were no condition differences: Alc vs AQ10: D(825)=0.0643, *p*=0.3539; Alc vs AQ60: D(840) = 0.0636, *p*=0.3558; AQ10 vs AQ60: D(771)=0.0548, *p*=0.5973. For PC3 (~3.9% of variance) there were no condition differences: Alc vs AQ10: D(825)=0.0474, *p*=0.7381; Alc vs AQ60: D(840)=0.1016, *p*=0.0248; AQ10 vs AQ60: D(771)=0.0984, *p*=0.0448; based on correcting for multiple comparisons (p<0.05/9 and thus p<0.0056 needed for significance). Right column shows abundance of PC values for each PC for each condition. Bar black: percent of neurons with PC values less than −0.01. Light gray bar: percent of neurons with PC values between −0.01 and +0.01. Dark gray bar: percent of neurons with PC value greater than +0.01. These add up to 100% of neurons for each drinking condition. There were no condition differences for PC1 or PC2 (PC1: X-squared = 1.7065, *p*=0.7895; PC2: X-squared = 6.1793, *p*=0.1862), while there was a significant difference in the distribution of PC values for PC3 (X-squared = 11.978, *p*=0.0175). This concurs with the data in previous paragraph suggesting that AQ60 was different from other drinking conditions in PC3, and with more higher and lower PC3 value neurons. Further work will be required to understand this difference, especially since PC3 explained <4% of total variance.

**Suppl.Fig.7.**
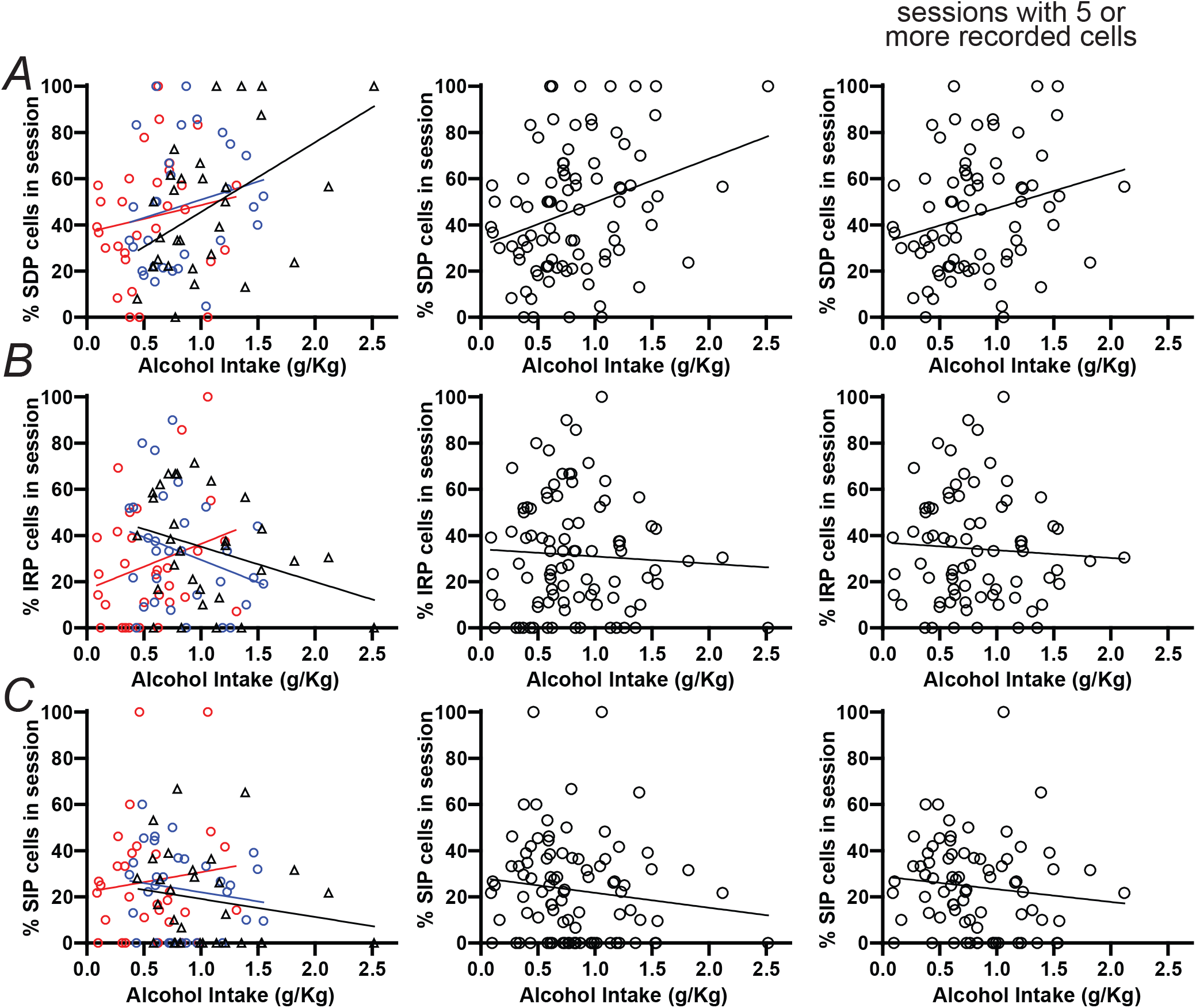
Abundance of cell phenotypes in relation to alcohol intake level. Left, middle, and right show (left) relation separated by drinking conditions, (mid) relation collapsed across drinking conditions, and (right) relation collapsed across drinking conditions and only sessions with 5 or more recorded cells (n=75). **(A)** g/kg level overall correlated with %SDP cells (*p*=0.0033), but, when separated by conditions, only for AOD (*p*=0.0059) (*p*>0.2 for AQ10, AQ60); *p*=0.0315 across 75 cells. **(B)** g/kg level did not relate to %IRP cells (*p*=0.5976 collapsed, *p*=0.0932 AOD, *p*=0.1343 AQ10, *p*=0.1637 AQ60; *p*=0.6104 for 75 sessions). **(C)** g/kg level did not relate to %SIP cells (*p*=0.2135 collapsed, *p*>0.3 for each condition alone; *p*=0.3076 for 75 sessions).

---

**Suppl. Table 1.** Correlation between total intake (g/kg alcohol) in a session and other session-level measures, including behavioral and in vivo firing measures. Significant relations are indicated in bold.

|  | P value | Slope | R<sup>2</sup> |
| --- | --- | --- | --- |
| *Behavior measures* |  |  |  |
| **Total Licks** | **<0.0001** | 298.5 | 0.465 |
| **Actual Lick Time** | **<0.0001** | 92.53 | 0.390 |
| **Avg Bout Length** | **0.0008** | 7.772 | 0.122 |
| Num of Bouts | 0.2518 | 1.654 | 0.015 |
| Avg IBI Length | 0.0627 | -0.807 | 0.040 |
| *Phenotype abundance* |  |  |  |
| **% SDP** | **0.0033** | 18.88 | 0.094 |
| % IRP | 0.5976 | -3.130 | 0.003 |
| %SIP | 0.2135 | -6.453 | 0.018 |
| *Phenotype Firing* |  |  |  |
| Change in SDP | 0.4667 | -0.543 | 0.006 |
| Change in IRP | 0.2136 | -1.748 | 0.021 |
| Change in SIP | 0.1515 | 1.799 | 0.033 |
| *Firing All cells in session* |  |  |  |
| Basal | 0.6347 | 0.329 | 0.003 |
| Lick Onset | 0.7662 | -0.353 | 0.001 |
| Macro Bout | 0.6223 | -0.469 | 0.003 |
| Actual Lick Time | 0.5815 | -0.541 | 0.003 |
| First Bout | 0.9019 | -0.120 | 0.000 |
| IBI | 0.0892 | -2.095 | 0.032 |
| Post-Lick | 0.9142 | -0.073 | 0.000 |
| *Basic ephys.* |  |  |  |
| # cells recorded | 0.6164 | 1.076 | 0.003 |
| Waveform width | 0.8015 | -0.055 | 0.001 |
| Waveform amplitude | 0.8374 | -2.008 | 0.000 |

